# Micromechanical Characterisation of 3D Bioprinted neural cell models using Brillouin Microscopy

**DOI:** 10.1101/2021.08.17.456575

**Authors:** Maryam Alsadat Rad, Hadi Mahmodi, Elysse C. Filipe, Thomas R. Cox, Irina Kabakova, Joanne L. Tipper

**Affiliations:** School of Biomedical Engineering, Faculty of Engineering and Information Technology, University of Technology, Sydney, NSW, Australia; School of Mathematical and Physical Sciences, University of Technology Sydney, Ultimo, NSW 2007, Australia; The Kinghorn Cancer Centre, Garvan Institute of Medical Research, Sydney, NSW 2010, Australia; St Vincents Clinical School UNSW Sydney, NSW, 2010, Australia

**Keywords:** 3D Bioprinting, GelMA hydrogels, Neuronal cells, Brillouin light scattering, Brillouin Microscopy

## Abstract

Biofabrication of artificial 3D in vitro neural cell models that closely mimic the central nervous system (CNS) is an emerging field of research with applications from fundamental biology to regenerative medicine, and far reaching benefits for the economy, healthcare and the ethical use of animals. The micromechanical properties of such models are an important factor dictating the success of modelling outcomes in relation to accurate reproduction of the processes in native tissues. Characterising the micromechanical properties of such models non-destructively and over a prolonged span of time, however, are key challenges. Brillouin microscopy (BM) could provide a solution to this problem since this technology is non-invasive, label-free and is capable of microscale 3D imaging. In this work, the viscoelasticity of 3D bioprinted neural cell models consisting of NG 108-15 neuronal cells and GelMA hydrogels of various concentrations were investigated using BM. We demonstrate changes in the micro- and macro-scale mechanical properties of these models over a 7 day period, in which the hydrogel component of the model are found to soften as the cells grow, multiply and form stiffer spheroid-type structures. These findings signify the necessity to resolve in microscopic detail the mechanics of in vitro 3D tissue models and suggest Brillouin microscopy to be a suitable technology to bridge this gap.

## 1. Introduction

Injury to the central nervous system (CNS) alters the molecular and cellular composition of neural tissue and leads to complex fibrotic networks of various stiffness, with scar-like tissue in particular inhibiting the regrowth of damaged axons [1]. The complexity of the fibrillary network makes it challenging to understand the reactions of different types of CNS cells, primarily neurons and glial cells, to mechanical injury such as spinal cord injury *in vivo* [2]. Due to the ethical issues in the use of animal models for tissue engineering (such as animal sacrifice), there is an increasing demand for accessible and valid (biologically relevant) in vitro neural tissue models which allow the user to track normal and aberrant cellular and molecular interactions, as well as whole system interconnectivity [3, 4].

Currently, tissue engineered biomaterials with specific mechanical properties which closely mimic the characteristics of the native CNS matrix are being developed. GelMA hydrogels are extremely promising candidates for use in such models, due to their tuneable physical properties and excellent biocompatibility [5, 6]. Determination of the mechanical properties of GelMA hydrogels as a key parameter to model native neural tissues has been investigated using several mechanical characterisation methods such as Atomic Force Microscopy (AFM) [7] or rheology [8]. The former measures the local quasi-static Young’s modulus (E) from the deflection of a cantilever. The latter, which is the most commonly used technique, allows characterisation of the average rheological properties in bulk materials. For example, the mechanical properties of acellular GelMA hydrogel samples with high, medium, and low methacrylation at GelMA concentrations of 5%, 10%, and 15% has been assessed using unconfined compression tests [9].

In the recent years, the majority of the studies have utilized rheology techniques to assess the mechanical properties of cell-laden GelMA hydrogels, which are used to fabricate scaffold-based constructs. For example, Yin *et al*. applied rheology to measure the mechanical properties of GelMA hydrogels with different polymer concentrations to obtain the optimized bioprinting setup [10]. In another study [11], rheological measurements were conducted for GelMA/gellan gum (GG) composite bioinks to investigate the effect of different cell densities on printing outcomes. The down side of conventional mechanical tests, either macroscopic techniques such as shear rheology or microscopic approaches such as AFM, is that they require mechanical forces to be directly exerted on the cellular hydrogels [12]. In addition, the other potential issue is that applying these forces activates intracellular force/mechanosensors, which may alter the cell behaviour and create artefacts in the system under study through repeated measurements [13]. These existing methods are difficult to extend to the measurement of cellular hydrogels, as relatively high contact forces can be damaging to the cells within the sample. In addition, rheology lacks microscale resolution [14] but AFM indentation can only be applied to measurements at the surface (2D) [13]. Ultrasound elastography is another imaging modality that is sensitive to tissue stiffness [15] but does not have microscale resolution because of relatively large irradiation area. Therefore, these techniques may not be suitable for real-time and continuous monitoring of bioprinted 3D cellular constructs.

Most recently, Brillouin microspectroscopy, a non-contact and label-free technique, has been demonstrated to be useful in characterising living organisms and artificial biomaterials, providing a unique insight into viscoelasticity at the microscale [16, 17]. The technique is based on Brillouin light scattering (BLS), which is an inelastic scattering process arising from the interaction of light with thermally driven acoustic phonons [18]. Unlike quasi-static techniques such as compression or shear rheology, Brillouin spectroscopy provides information on the complex longitudinal elastic modulus of the material in a high-frequency regime (∼1-10 GHz) [19]. The 3D micromechanical mapping of samples can be achieved using a combination of confocal microscopy with Brillouin spectroscopy, known as Brillouin microscopy (BM) [20]. In BM, the mechanical mapping is achieved by scanning the sample with a low-power focused laser beam, thus making this technology compatible with *in situ* and *in vivo* imaging in 3D. This technique has been applied to investigate the mechanical properties of the corneas [21], tumours [22], fibrous proteins of the extracellular matrix [23, 24], cellular mechanics [25, 26], and more recently, the first mechanical images of a mouse embryo [27, 28]. In addition to the studies of living cells and tissues, Brillouin spectroscopy has been employed to characterise the microscopic viscoelastic properties of various bio-compatible materials, including casted and bioprinted hydrogels [29, 30]. Overall, the non-contact and label-free nature of BM has helped this technology grow in popularity, in particular for application in *in vivo* settings for biomedical diagnostics and disease monitoring [16].

In this work, GelMA hydrogel constructs were bioprinted in an extrusion-based device, and the microscale viscoelastic properties of the 3D bioprinted hydrogel models were assessed for the first time by using BM. The mechanical properties of hydrogels were altered by varying the polymer concentration and the degree of cross-linking with UV exposure. Here, we show that the Brillouin frequency shift of hydrogels is more sensitive to polymer concentration than to the effects of UV exposure time. We validate Brillouin microscopy results for GelMA hydrogels using industry-standard rheology measurements. The effects of incorporating NG 108-15 neuronal cells into the GelMA hydrogels were then investigated at both macro- and micro-scale over 7 days. We found that neuronal cells have high viability and high proliferation rates in low polymer concentration hydrogels (2.5 and 5% (w/v)). At the macroscale, Brillouin frequency shifts in cellular hydrogels was found to be significantly lower than that of acellular hydrogels at any measurement time. At the microscale, the neuronal cells exhibited spheroid morphology and on average showed significantly higher Brillouin frequency shifts than that of the surrounding hydrogel, with difference growing over 7 days. These results demonstrate that the mechanical properties of acellular and cellular hydrogels are time-dependent rather than static, with evolution depending on the cellular growth and metabolic activity, gel degradation and external incubation factors. The details of the time evolution of 3D bioprinted models need to be considered for designing a valid model and correct interpretation of biological results. In this work we have shown that such details are possible to obtain by using non-destructive technologies compatible with 3D label free imaging, such as BM.

## 2. Methods

### 2.1. Preparation of GelMA hydrogels with different volume concentrations

The CELLINK GelMA Kit (Cellink, Sweden) containing lyophilised Gelatin methacryloyl (GelMA) and Lithium acylphosphinate photoinitiator (LAP) was used in this study. Initially, LAP powder was mixed with Phosphate-buffered saline (PBS; GIBCO, Thermo Fisher Scientific) (1X, pH 7.4) for 20 min at 60 °C to prepare a LAP solution with a concentration of 0.25% (w/v). After the mixture was dissolved entirely and was homogenous, the solution (PBS/LAP) was filter sterilised (0.22 μm) in a Class II biosafety cabinet and wrapped in aluminium foil to protect it from light. The desired volume of the sterilised photoinitiator was added to GelMA to achieve the hydrogel concentrations with ranges from 2.5 to 15% (w/v). For example, for making 10 % (w/v) GelMA with 0.25% (w/v) LAP concentration, 5 ml of photoinitiator was added to 500 mg GelMA powder with stirring and heating at 50°C. Finally, GelMA hydrogel was transferred into a 3 mL Luer-lock syringe (CELLINK) for bioprinting. For printing cellular hydrogels, NG 108-15 neuronal cells at a concentration of 6 × 10^6^ cells.mL^-1^ were mixed with GelMA hydrogels with different concentrations, 2.5, 5, 7.5, 10, and 15% (w/v) at a ratio of 1:10.

### 2.2. Three dimensional (3D) Bioprinting of hydrogels

Bioprinting of GelMA hydrogel was performed using a BIO X 3D Bioprinter (CELLINK, Gothenburg, Sweden). Lattice scaffolds with dimensions of 3×3×1 mm^3^ (width× length× height), a grid size of 1.05 mm, and 45% porosity were selected as a printing model. Prior to printing, the bioink solution (GelMA hydrogel) was heated to 37 °C in an incubator and then was loaded into the bioprinter system, allowed to cool to the desired temperature, between 22-33°C, based on the gelation point of the different GelMA hydrogels concentrations of 2.5 to 15 % (w/v). The bioinks with different concentrations were printed with a 22G nozzle size at optimised pressures between 9 and 35 kPa and speeds of 5-11 mm/s. The bed temperature of the bioprinter was kept at 10°C to allow thermal gelation of GelMA bio-inks during printing (prior to UV cross-linking). The bioprinted GelMA hydrogels were subsequently cross-linked by UV exposure (405 nm at an intensity of 19.42 mW.cm^-2^) for between 30 and 300 s. Three printed samples for each concentration were immersed in DMEM cell culture medium cell culture media containing 10 % (v/v) FBS and incubated at 37°C, 5% (v/v) CO_2_ in the air to reach the equilibrium swelling state before measurements were taken.

### 2.3. NG 108-15 neuronal cell culture

The NG108-15 (mouse neuroblastoma × rat glioma hybrid, HPA Culture Collections) cell line was purchased from the European Collection of Cell Cultures (ECACC). This cell line was grown in DMEM containing 10% (v/v) FBS (Gibco), 1% (v/v) penicillin-streptomycin (10000□Units.ml^-1^-10000 μg.ml^-1^) (Gibco), and 2% (v/v) HAT (0.1 mM hypoxanthine, 400 nM aminopterin and 0.016 mM thymidine, Sigma Aldrich). When cells were approximately 80% confluent, the media was removed, and the cells were washed with 10mL Dulbecco’s phosphate-buffered saline (DPBS) (Gibco; without calcium and magnesium) to remove any remaining media. Then, the DPBS was replaced with 2 mL of a 0.25 % (v/v) trypsin/EDTA (Sigma-Aldrich), and the cells were incubated at 37° C for 5 min at room temperature. The cell suspension was transferred into a sterile universal tube containing 10mL cell culture media containing 10 % (v/v) FBS to neutralise the trypsin and centrifuged at 400 g for 5 minutes. The waste media containing trypsin was removed, and the pellet of cells was re-suspended in 2 ml of the appropriate cell culture media before mixing with hydrogels.

### 2.4. Determination of cell viability using live/dead assay

Cellular hydrogels were rinsed with PBS and stained with 2 μg.ml^-1^ Calcein AM (Sigma-Aldrich), 1 μg.ml^-1^ propidium iodide (Sigma-Aldrich), and 2 μg.ml^-1^ Hoechst (Thermo Fisher) for 30 min at 37°C in 5% (v/v) CO_2_ in air. The staining solution was discarded, and hydrogels were washed four times with DPBS, and images of cellular hydrogels were acquired using an inverted fluorescence microscope (Olympus IX 73 & CKX3).

### 2.5. Determination of cell viability using ATP assay

The viability of NG108-15 neuronal cells was measured via Luminescent Cell Viability Assay on days 1, 3 and 7. The assay procedure involved adding a single reagent directly to cells, resulting in cell lysis and generation of a luminescent signal proportional to the amount of ATP present. In this research, Celltiter-Glo^®^ reagent (Promega) was added to each of the cellular GelMA hydrogel samples according to the manufacturer’s protocol. The cell culture medium was removed entirely and replaced with the ATP reagent at a volume that fully immersed the hydrogels. This process was followed by incubating cellular hydrogel samples on a shaker for 45 min. A microplate reader (Varioskan™ LUX, Thermo Scientific) was used to record luminescence (counts per second) for each well.

### 2.6. Biomechanical Characterisation of hydrogels

The Young’s modulus of acellular GelMA hydrogel samples (*n* = 3) was obtained through unconfined compression and shear rheology on a TA Instruments Dynamic Hybrid Rheometer (DHR-3, TA Instruments). An 8mm diameter punch biopsy was taken from each hydrogel and placed between the top and bottom parallel plate geometries. Mechanical characterisation of the hydrogels was performed using shear rheology and unconfined compression tests, with a constant compression rate of 10 mm/min with data output in the form of axial force (N) and gap (mm). For unconfined compression testing, the data was analysed and a stress/strain curve for each replicate hydrogel was obtained. The Young’s modulus (kPa) was obtained from the linear region of the stress/strain curve. For shear rheology, amplitude sweeps were performed at frequencies of 0.159 HZ for strain amplitudes ranging from 0.2 % to 2%. This determined the shear strain range within the Linear Viscoelastic Region (LVER) before deformation of the material occurred. All samples were measured at 25° C and on the same day and time with Confocal Brillouin Microscopy to have the same swelling conditions.

### 2.7. Confocal Brillouin Microscopy

#### 2.7.1 Brillouin Characterization

The GHz frequency viscoelastic properties of cellular and acellular GelMA hydrogel samples were also assessed using BM. For acellular GelMA hydrogels, prior to rheological characterisation, the same hydrogels were analysed using BM. This system was used to collect the spontaneous Brillouin scattering spectra and the system comprised of a single-frequency solid-state laser (Torus, 660 nm), a confocal microscope (CM1, JRS Instruments), a 3D scanning microscopy stage (SmartAct) and a tandem Fabry-Perot interferometer (TFP1, JRS Instruments). The spectrometer resolution was approximately 360 MHz, and the spectral extinction ratio was above 10^10^. We used objective lenses of 20X (Mitutoyo, NA=0.42, WD=20 mm) and 60X (Nikon, WD=2.8 mm, NA=1) for macro- and micro-mechanical characterisation of hydrogels, respectively. The axial resolution of the Brillouin system was approximately 120 µm and 12 µm for the objective lens of 20X and 60X, respectively, by considering 300 and 100 µm input aperture sizes in front of the spectrometer. The lateral resolution (XY plane) for our system was estimated to be below 2 μm for both objectives. The laser power for acellular hydrogel samples was 100 mW, and for cellular hydrogel samples this was kept at less than 20 mW to avoid sample damage by incident radiation. A schematic of the BM is illustrated in Fig. 1A.

**Fig 1.**
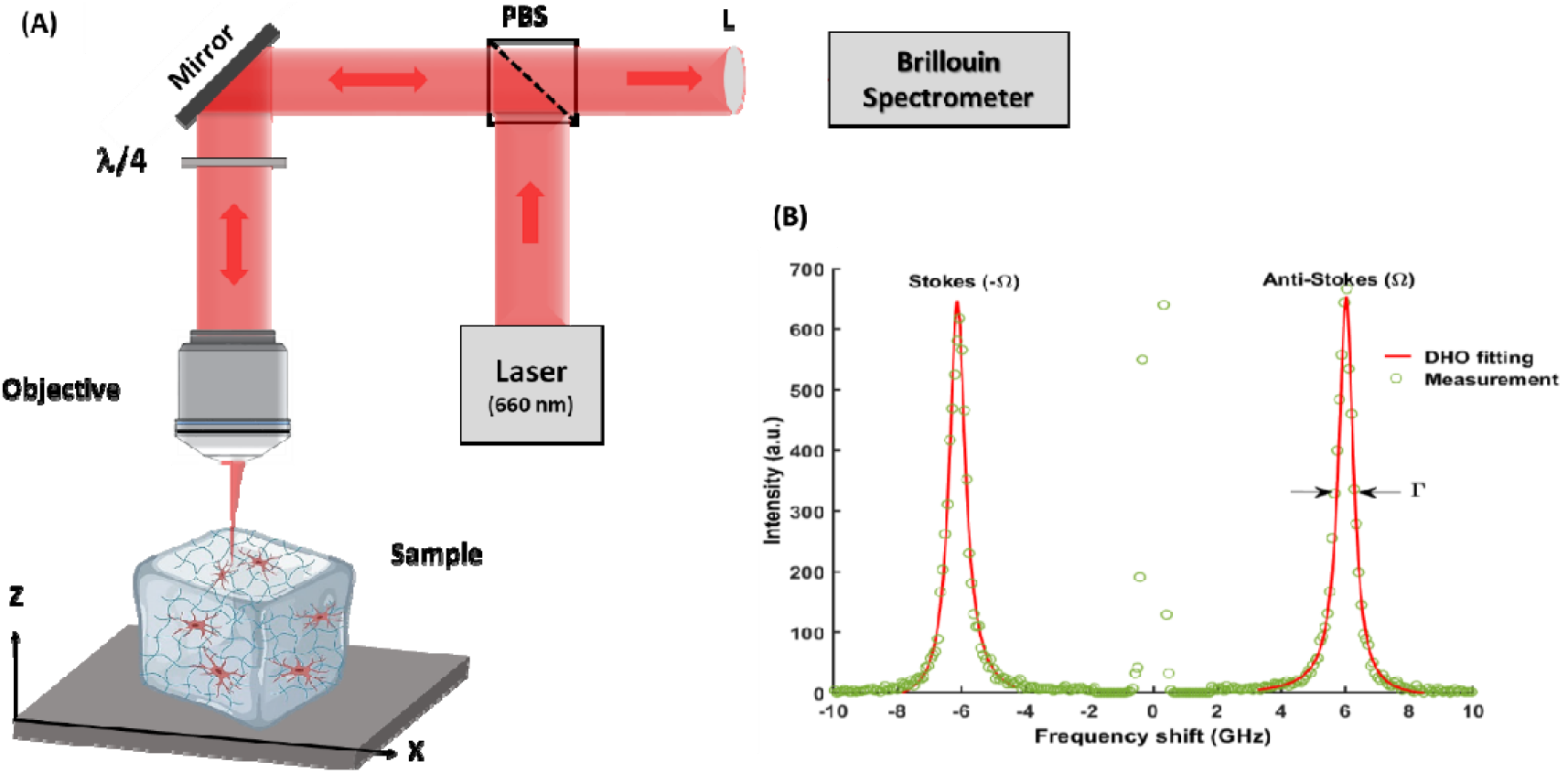
(A) Schematic of Brillouin microscope (BM). Notations used: L-lens, PBS-polarisation beam splitter, λ/4 - quarter-wave plate. (B) A typical Brillouin spectrum showing the Stokes and Anti-Stokes peaks.

A spontaneous Brillouin light scattering spectrum consists of two signals located at lower (Stokes, -Ω) and upper (Anti-Stokes, Ω) Brillouin frequency with respect to the incident light frequency (for simplicity adjusted to be at 0 GHz) as shown in Fig. 1B. Two independent parameters can be extracted from the Brillouin spectrum: Brillouin frequency shift (BFS) and Brillouin linewidth (*Γ*), which are determined from the exact position and linewidth of Stokes and Anti-Stokes peaks, respectively. The frequency shift of Brillouin peaks can be presented as [31]:

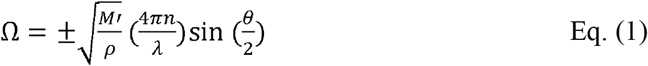

where *M*′ is the longitudinal storage modulus, *ρ* is the mass density, *n* is the refraction index, *λ* is the incident wavelength, and *θ* is the scattering angle, which is equal to 180° (backscattering) in the utilised optical setup. Moreover, the Brillouin linewidth is expressed by

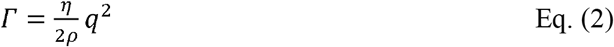

Where *η* is proportional to the material viscosity and 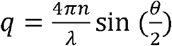 is the wave vector. Based on the equations (1) and (2), the complex longitudinal viscoelastic modulus (*M\**), which is composed of elastic storage (*M′*) and loss (*M′′*) moduli are given by

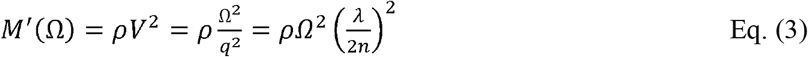

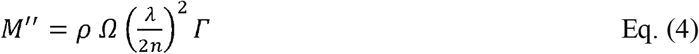

#### 2.7.1. Brillouin data collection and processing

Brillouin microscopy data was collected using commercial Ghost (JRS Instruments) and in-house software. The collected data was acquired from single point measurements and line scanning, which was performed by scanning the sample plate in the X, Y or Z directions on the microscopy stage while keeping the optical system fixed. The Brillouin frequency shift and the linewidth of the hydrogel samples were obtained at three random locations for each sample and the results were averaged. The Brillouin frequency shift and the linewidth were determined by fitting with a damped harmonic oscillator (DHO) model to every Stokes and anti-stokes Brillouin peak. The acquisition time for each point measurement was 20 seconds to optimise signal-to-noise-ratio and improve the fitting precision. In addition, a deconvolution of the instrumental function response was applied to correct for the finite linewidth of the laser spectrum (Fig. S1, see Supplementary information for details).

### 2.8. Statistical analysis

Calculated data points are presented as mean ± standard deviation. One-way and two-way ANOVA followed by Tukey post-hoc correction tests were performed where appropriate to measure statistical significance (GraphPad Prism 5.02 Software). Differences were taken to be significant at p < 0.05.

## 3. Results

### 3.1 Acellular GelMA hydrogel characterisation

#### 3.1.1 Effect of UV cross-linking on stiffness

The acellular GelMA hydrogels were printed using optimised parameters which are given in Table S1 and Fig. S2 in the supplementary data. The effect of UV cross-linking on viscoelasticity of bioprinted acellular GelMA hydrogels in GHz frequency range was investigated using BM. As shown in Fig. 2, a significant difference was observed in the BFS values between the UV cross-linked samples for 120 s and those not exposed to UV light for gels of all solid fractions. However, this difference in BFS was found to be more significant for the concentration equal to or above 5% (w/v) (P< 0.0001) compared to 2.5% (w/v) (p < 0.05). The results presented in Fig. 2 demonstrate that, by increasing the solid fraction of the GelMA hydrogels, the UV exposure time required to effectively crosslink the samples decreased.

**Fig. 2.**
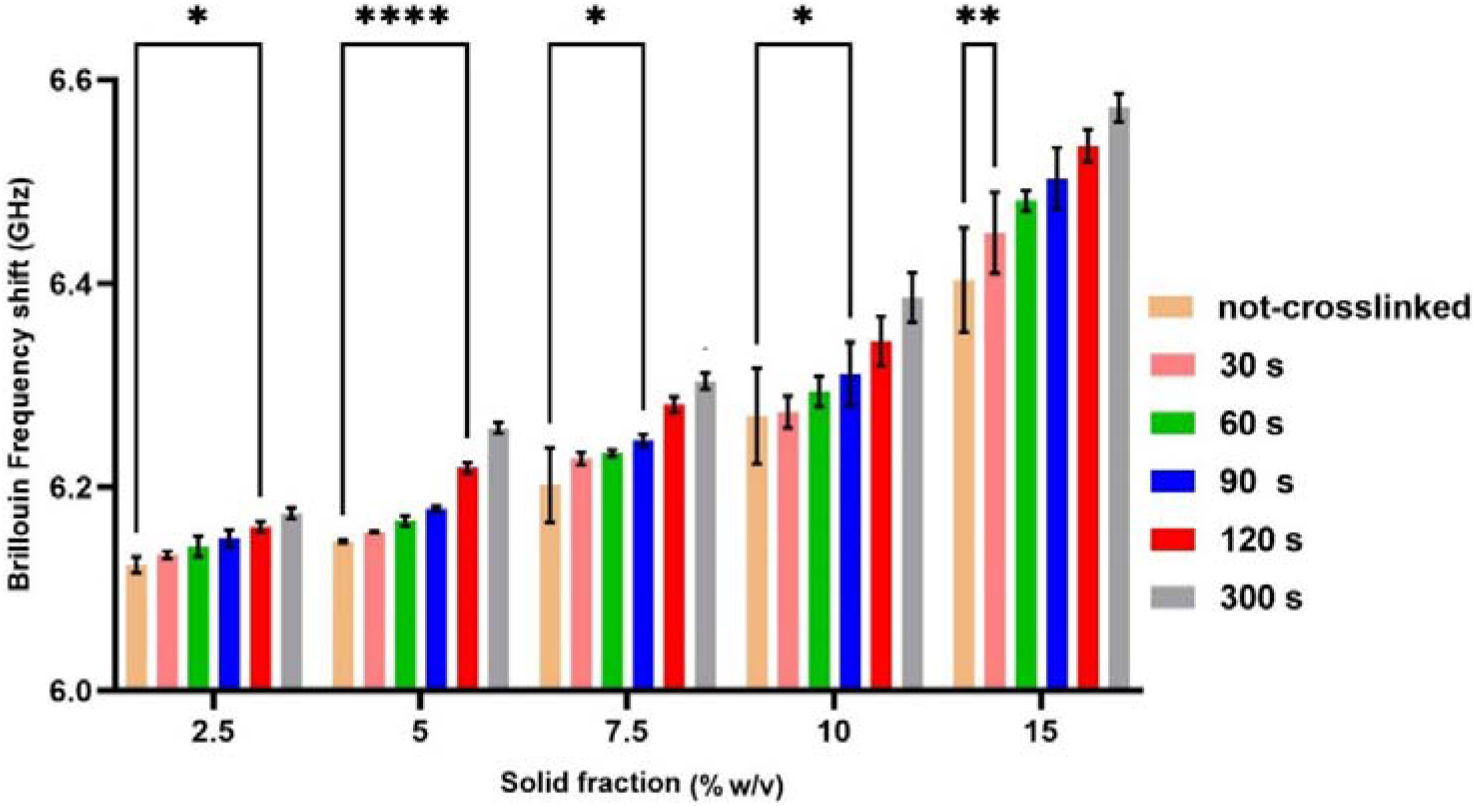
Brillouin frequency shift of bioprinted acellular GelMA hydrogels with different concentrations of 2.5, 5, 7.5, 10 and 15 % (w/v) and LAP concentrations of 0.25 % (w/v). The printed hydrogels were cross-linked under UV light with wavelength of 405 nm and an intensity of 19.42 mW.cm^-2^ from 30s to 300s. Results are given as mean ±SD (n = 3). Two-way ANOVA followed by Tukey post-hoc correction tests, significance levels: *P < 0.05, **P P <0.001, ****P <0.0001.

#### 3.1.2 Effect of solid fraction variation on stiffness

The effect of solid fraction variation on viscoelasticity of the acellular GelMA hydrogels was investigated. We found that both BFS and linewidth increased with increasing solid fraction, with BFS showing significant differences for all concentrations (****P <0.0001, not shown in Fig. 3a). In contrast, linewidth values exhibited a significant difference only between two concentration pairs 7.5%-10 % (w/v), and 10%-15% (w/v), as displayed in Fig. 3b. The results show a positive correlation between the Brillouin frequency shift/linewidth and the solid fraction. Based on BFS and linewidth values, the storage (*M’*) and loss (*M″)* longitudinal moduli were also calculated and plotted in Fig. 3a and 3b. Since it is challenging to directly measure the refractive index and density in bulk samples, the values for *M’* and *M″* reported here should be treated as estimates only. To do so, we assumed a constant refractive index of *n*=1.35 and a density of *ρ*=1030 kg/m^3^, based on previous studies [32].

**Fig. 3:**
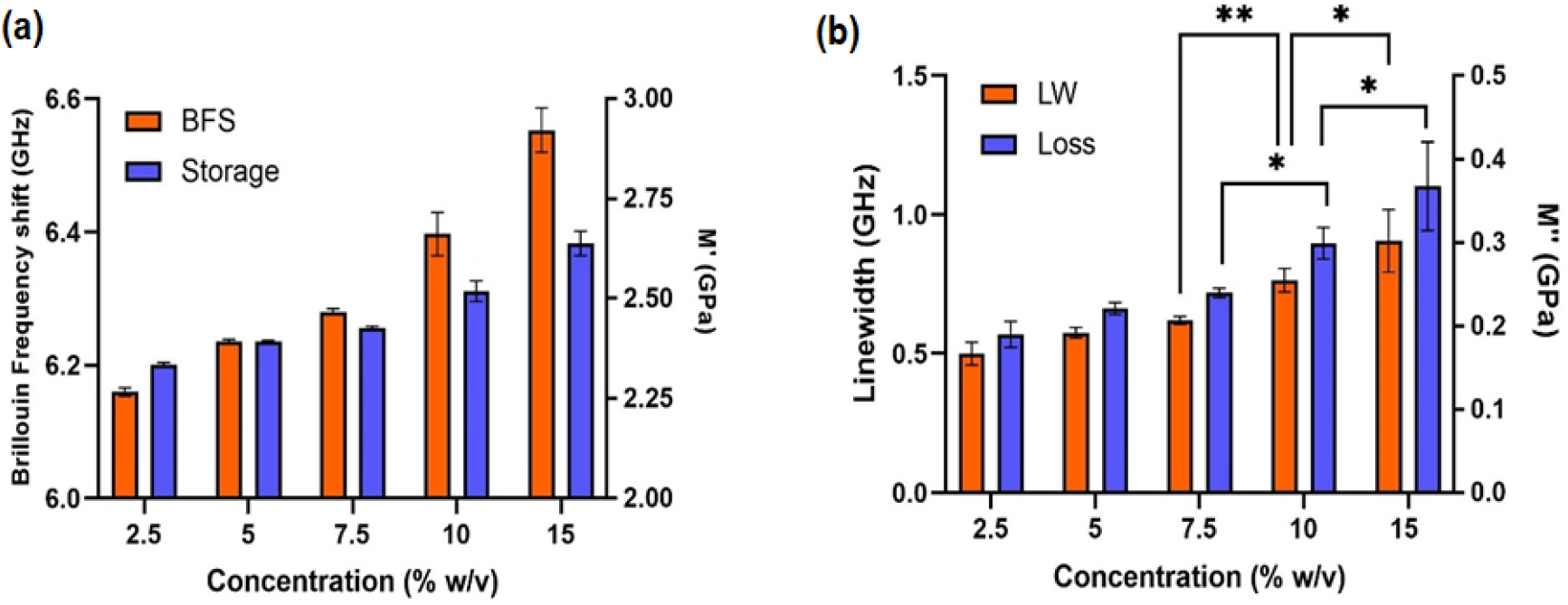
(a) Brillouin frequency shift and calculated storage longitudinal modulus and (b) linewidth and calculated loss longitudinal modulus for 3D bioprinted acellular GelMA hydrogels of different concentrations. Results are given as mean ±SD (n = 3). Two-way ANOVA, significance levels: *P< 0.05, **P< 0.001.

The viscoelastic properties of bioprinted acellular GelMA hydrogels were also investigated using unconfined compression and rheology to verify the accuracy of data achieved using Brillouin Microscopy. The Young’s and shear moduli of samples were plotted against hydrogel solid fractions (Fig. S3). Increasing the volume concentration of GelMA in the hydrogels from 2.5% to 15% (w/v) increased the compressive modulus by almost 14-fold, as shown in Fig. S3 (b)). To assess the correlation between BM and rheology measurements, storage longitudinal modulus (*M′*) was plotted against storage shear modulus (*G*′) for all GelMA hydrogel concentrations. As shown in Fig. 4, a six-fold increase in solid fraction resulted in a thirty-five-fold increase in storage shear modulus, where storage longitudinal modulus was increased by 14%.

**Fig 4.**
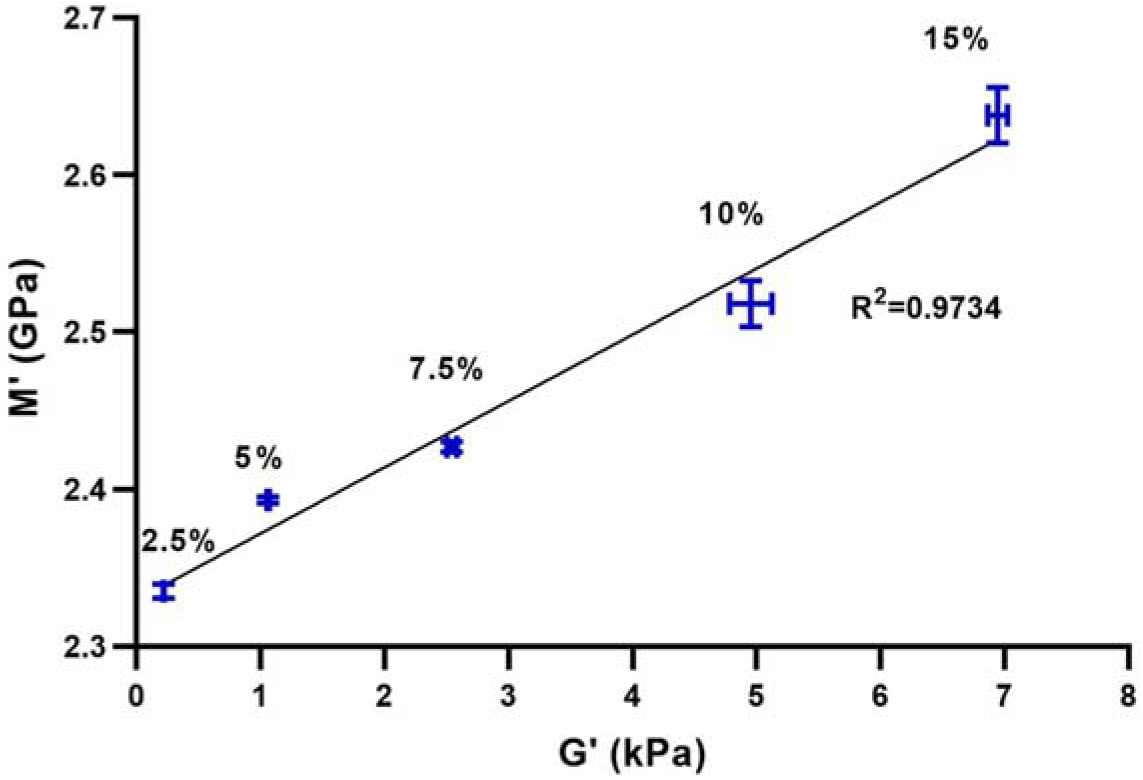
Correlation between storage longitudinal (*M′*) and shear moduli (*G′*) for bioprinted acellular GelMA hydrogels with concentrations of 2.5, 5, 7.5, 10 and 15% (w/v) measured using BM and rheology. Results are given as mean ±SD (n = 3). The solid line shows a linear regression across the measured data with R^2^ = 0.9734.

### 3.2 Cellular GelMA hydrogel characterisation

#### 3.2.1 Cell viability

Bright-field and fluorescent microscopy images of NG108-15 neuronal cells grown in different GelMA hydrogels with concentrations of 2.5, 5, 7.5, 10% (w/v) are displayed in Fig. 5. The lowest NG108-15 cell viability was observed for the GelMA hydrogel with a concentration of 10 % (w/v). In comparison, higher cell viability and lower numbers of dead cells were observed for GelMA hydrogels with concentrations of 2.5 and 5 % (w/v) (Fig. 5 a-d). Furthermore, no cell elongation was observed in the morphology of NG 108-15 cells in the GelMA hydrogels network. However, the cells proliferated and organised into spheroid/aggregates in concentrations of 2.5 and 5 % (w/v) over the seven days.

**Fig 5.**
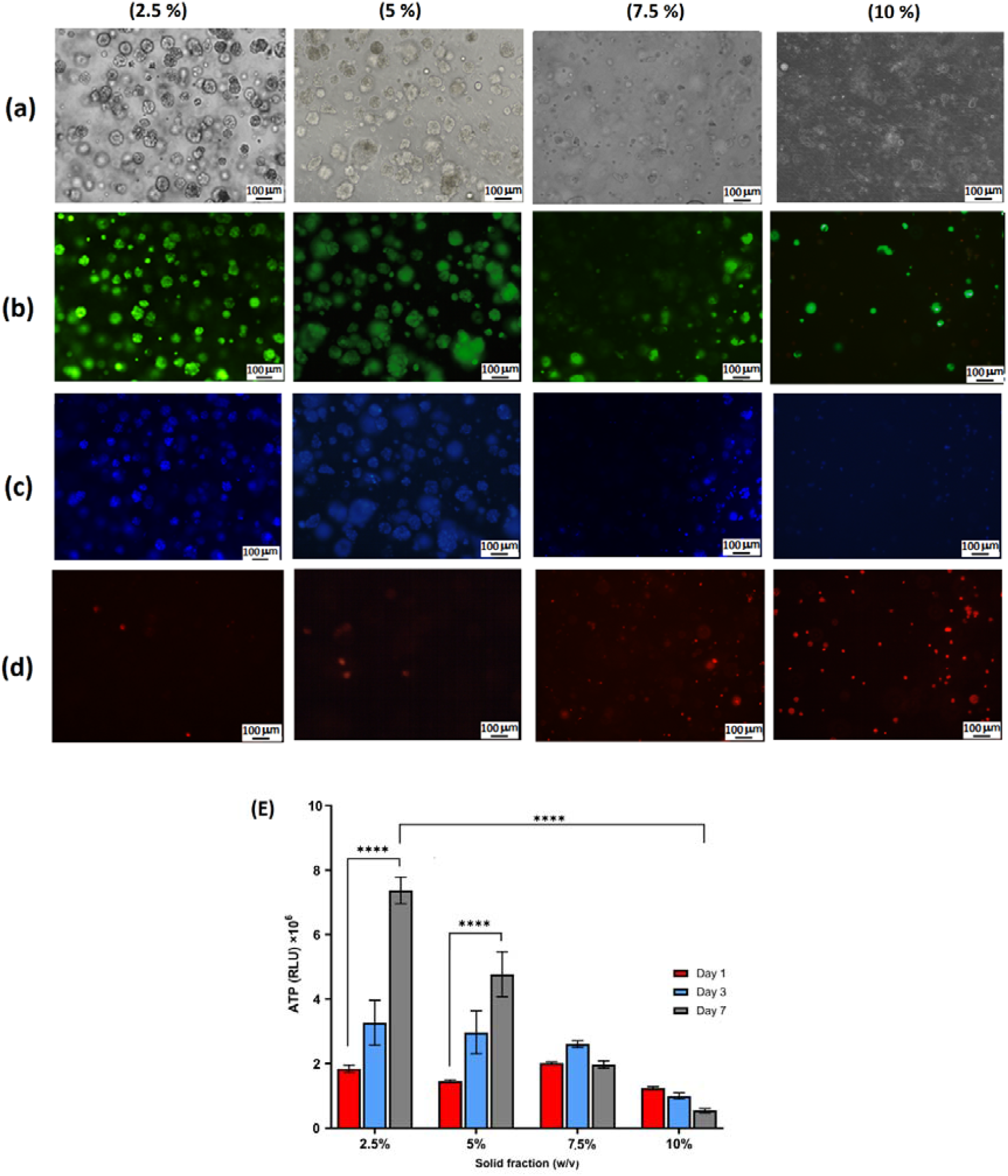
Representative bright-field (a) and fluorescence microscopy (b-d) images of NG108-15 neuronal cells in different GelMA hydrogels with concentrations of 2.5-10 % (w/v) after 7 days at 37°C, 5% (v/v) CO_2_ in air. The cytoplasm of live cells was stained with Calcein AM and is shown in green, and the nuclei stained with Hoescht appear blue. Dead cells were stained with propidium iodide (PI) and are stained red. (e) Quantitative cell viability assay results. Error bars indicate 95% confidence intervals; lines indicate statistical diCerences determined by two-way ANOVA, n = 3. Significance level: ****P<0.0001.

In addition, NG 108-15 cell viability was determined by quantifying the ATP content of cells on days 1, 3 and 7 (Fig. 5e). The number of viable NG108-15 cells for GelMA hydrogels with concentrations of 2.5 % and 5% (w/v) was found to be significantly higher on day seven compared to the first day. However, no significant di□erences were observed for cell viability in GelMA hydrogels with concentrations of 7.5 % (w/v) between the first and third days. Moreover, the ATP content was decreased for cellular GelMA hydrogels with a concentration of 10% (w/v) over seven days. Overall, the qualitative and quantitative cell viability data showed that NG108-15 neuronal cells have optimal proliferation rates and lower numbers of dead cells in the 2.5% (w/v) GelMA hydrogels relative to ones with higher GelMA volume concentrations.

#### 3.2.2 Macromechanical properties

The influence of NG108-15 neuronal cell incorporation on the average macromechanical properties of GelMA hydrogels was investigated by taking Brillouin point measurements with a low NA objective lens (20x) in random locations across hydrogel samples. For this purpose, three samples for each concentration of cellular hydrogels were prepared, as described in the methods section. The BFS of cellular GelMA hydrogels was measured and compared with acellular GelMA hydrogels, with individual measurements taking place on days 1, 3, and 7. Analysis showed that acellular GelMA hydrogels with concentrations of 2.5 %, 5% and 7.5 % had significantly higher BFS than cellular GelMA hydrogels on all three measurement days (Fig. 6a-c). However, assessment difference between 10% (w/v) cellular and acellular GelMA hydrogels was measured only on day 1 of the measurements, as shown in Fig. 6d.

**Fig 6.**
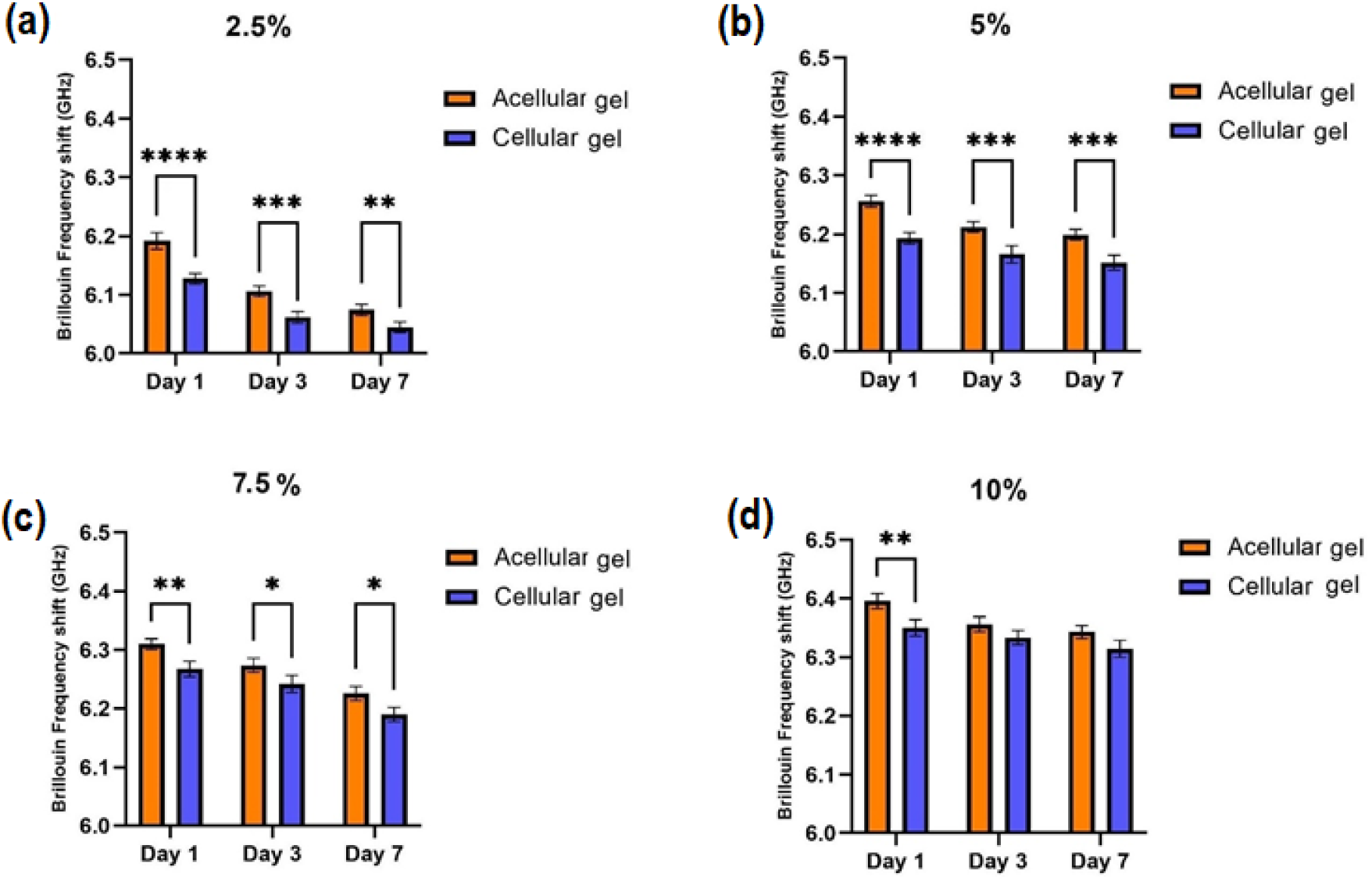
BFS measurements of the across the bioprinted acellular and cellular GelMA hydrogels embedded with NG108-15 neuronal cells at a concentration of 4 × 10^6^ cells/mL and concentrations of (a) 2.5%, (b) 5 %, (c) 7.5 % and (d)10 % (w/v). Error bars indicate 95% confidence interval; lines indicate statistical difference determined by two way ANOVA. Significance levels: *p < 0.05, **P <0.005, ***P <0.0005, ****P <0.0001, n = 3.

#### 3.2.3 Local micromechanical properties

The scanning of cellular GelMA hydrogels was carried out on days 1, 3, and 7 to study the dynamic micromechanical properties of cells within 3D extracellular hydrogel networks. For this purpose, only the cellular GelMA hydrogels, which had given the optimal cell viability results (2.5% (w/v)) using the quantitative cell viability assay (see Section 3.2.1), were analysed. Before the Brillouin measurement, the viability of neuronal cells was investigated using a live/dead assay at the same time point. The bright-field and fluorescent images of NG 108-15 cells showed high cell viability and the incremental increase in spheroid size over seven days (Fig. 7). The red line in the middle panel of Fig. 8 shows the laser scanning path with a step size of 1 µm. The average and standard deviation for the BFS were calculated based on averaging two measurement repeats for each sample. As shown in Fig. 8, the high and low-frequency shift values are attributed to the NG 108-15 neuronal cell spheroid and hydrogel, respectively. The BFS value of the NG 108-15 neuronal cell spheroid was 6.25±0.029, 6.30±0.034, and 6.43±0.039 GHz on day 1, 3 and 7 days, respectively. An apparent increase in BFS was observed over seven days, possibly due to the increasing spheroid size and number of cells, as is illustrated in the bottom panel of Fig. 8. The BFS values of cellular GelMA hydrogels were 6.13±0.019, 6.09±0.024, and 6.055±0.023 GHz on the 1, 3 and 7 days, respectively. In comparison, the average BFS values of acellular GelMA hydrogels were 6.20±0.010, 6.11±0.018, and 6.089±0.015 GHz on days 1, 3 and 7, respectively (Fig. 8a), which are higher than the BFS of cellular GelMA hydrogel but lower than that of cell spheroids.

**Fig. 7.**
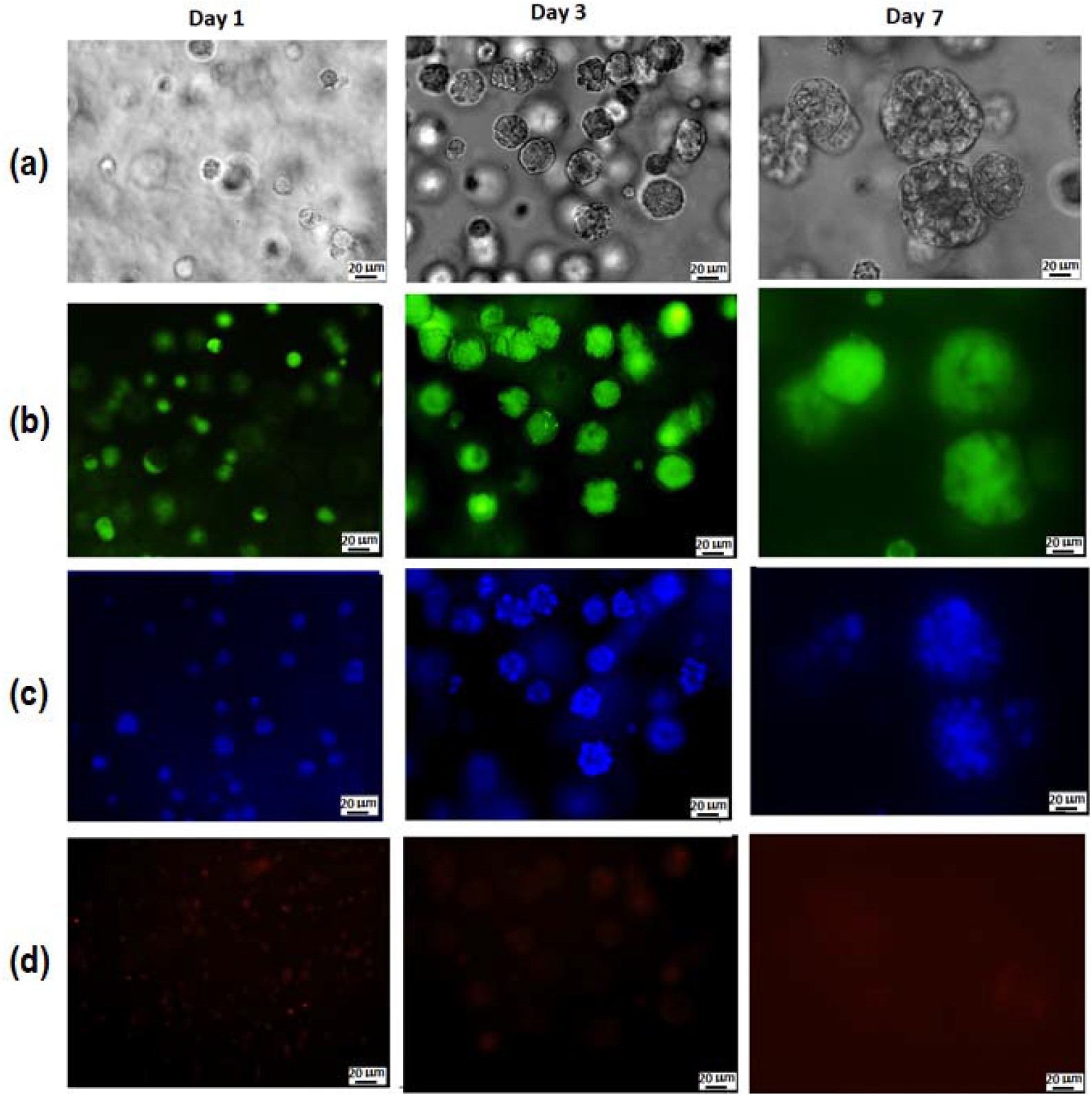
Representative bright-field (a) and fluorescence microscopy (b-d) images of NG108-15 neuronal cells embedded in a GelMA hydrogel with concentrations of 2.5% (w/v) at days 1, 3 and 7 at 37°C, 5% (v/v) CO_2_ in the air. The cytoplasm of live cells was stained with Calcein AM and is shown in green (b), and the nuclei stained with Hoechst were stained blue (c). Dead cells were stained with propidium iodide and are stained red (d).

**Fig. 8.**
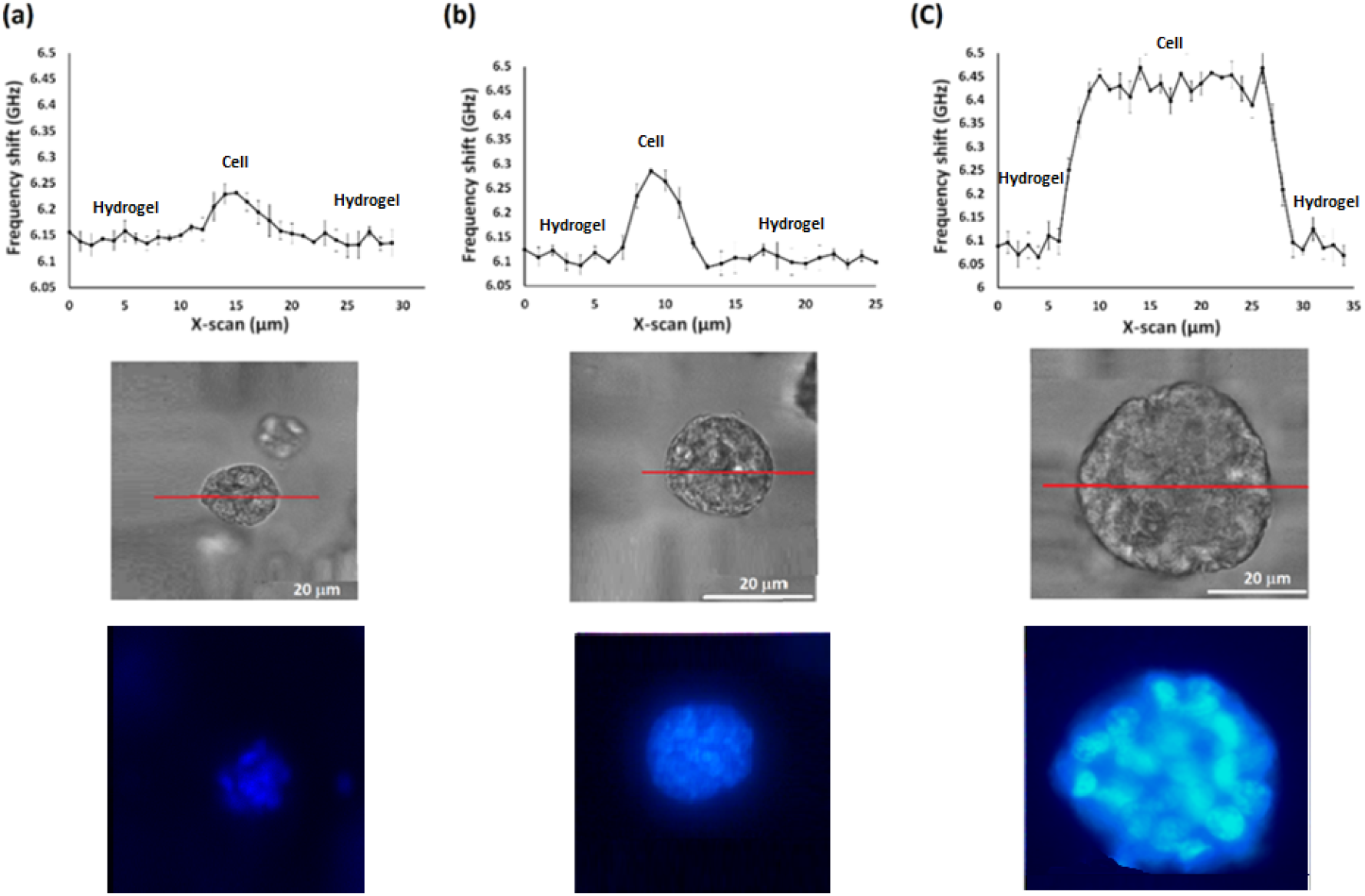
Brillouin line scanning (top panel), phase-contrast (middle panel), and fluorescent microscopy images (bottom panel) of neuronal cells within GelMA hydrogels with 2.5 % (w/v) concentration on days (a) 1, (b) 3, and (c) 7. The average and S.D are obtained based on averaging three measurement repeats for each hydrogel sample.

## 4. Discussion

BM was successfully utilised to investigate the mechanical properties of bioprinted acellular and cellular hydrogel constructs in 3D. Gel viscoelasticity was shown to alter as the result of solid fraction variation in the gel composition. These results are in full agreement with a previous study that reported increasing Brillouin frequency shifts with increasing polymer concentrations of GelMA hydrogel [19]. We also confirmed that the Brillouin linewidth/loss modulus of acellular GelMA hydrogels was more sensitive to changes in the gel’s solid fraction than the BFS/storage modulus, showing variation of 80.9% versus 6.5%, respectively, comparable to previous reports [29]. As the Brillouin linewidth is associated with the microscopic viscosity of the hydrogel, its variation could be attributed to the restricted mobility of water in hydrogels with high solid fraction, and consequently dense polymer networks [29].

The correlation between the storage longitudinal modulus and storage shear modulus showed a positive trend with respect to the solid fraction of hydrogels. A six-fold increase in solid fraction resulted in a thirty-five-fold increase in storage shear modulus, where storage longitudinal modulus was increased by 14%. This difference between the two measurements originates from the different physical meaning of these moduli, which assess different types of deformation. Brillouin spectroscopy probes the materials longitudinal modulus (inverse compressibility), which is a measure of stiffness under constrained deformation [33]. Rheology measurements, however, probe the material deformation under shear stress, with the material being unconfined in the measurement chamber. For the hydrogels with low solid fractions (<10%), the gel compressibility was shown to be influenced by the liquid phase of the aggregate material and showed significantly less variation [33].

The viability, morphology, and proliferation rate of NG 108-15 cells exhibited dependence on the solid fraction of the GelMA hydrogel. From the fluorescent microscopy images of the 3D cultured samples, the cells in softer hydrogels (2.5% and 5% (w/v)) were rounded and had a spheroid shape, which may have been due to the decreased numbers of arginine-glycine-aspartic acid (RGD) sequences in the lower hydrogel concentrations. It has been demonstrated that the addition of RGD (an integrin ligand) into the hydrogel networks can lead to better cell adhesion, proliferation, migration, and matrix production [34]. The quantitative cell viability results in the present study also showed that the cells in the stiffer hydrogels (7.5 % and 10 % (w/v)) were minimally proliferative, compared to the cells in the softer hydrogels (2.5 % and 5 % (w/v)). Banerjee and co-workers have shown that the proliferation of neuronal stem cells decreased as hydrogel stiffness increased [35]. Possible reasons for the low cell proliferation in hydrogels with high concentrations are low hydrophilicity and wettability of hydrogels. Previous studies of GelMA hydrogels, where concentrations was varied from 5 to 15 % (w/v), showed that the 5% GelMA hydrogel had the highest hydrophilicity and wettability, thereby facilitating the penetration of nutrients and oxygen [36].

Dynamic changes in the micromechanical properties of cellular and acellular hydrogel models over 7 days were probed by BM to determine the effect of cell incorporation within the 3D hydrogel matrix. The results indicated that the BFS of the hydrogel component of both cellular and acellular models with concentrations of 2.5 and 5 % (w/v) decreased over the 7 days, with cellular hydrogel models showing larger decreases in BFS than the acellular gels. A possible explanation of this finding could be cellular remodelling of the hydrogel matrix and therefore the stiffness of the cellular gels change over time [37]. It is noteworthy that the BFS of the cellular GelMA hydrogels with 10 % (w/v) concentration was significantly lower than acellular hydrogels only on the first day of the experiment. From the fluorescent microscopy images of the 3D cultured samples, we learned that the cells in stiffer matrices (7.5% and 10% (w/v)) were minimally adhesive, had low proliferative potential and high death rate over the 7 day experiment. These findings indicate that the low cell survival rates in the stiffer hydrogels limited the opportunity for the cell-matrix interaction and therefore matrix remodelling was minimal.

Brillouin microscopy measurement of cellular spheroids in this work was achieved using line scanning rather than 2D/3D mapping due to the limited speed of the TFP interferometer (10-20 s per single spectrum). In future studies, the acquisition time per point could be reduced down to 20-100 ms, with different instrument design, such as incorporating a virtually imaged phased array (VIPA) spectrometer [38] or by simulated Brillouin scattering (SBS) microscopy [39]. Overall, the time required to obtain a 3D image using a Brillouin microscope is a major challenge for this technology. Several improvements have been suggested, e.g. line-scanning microscopy [40], that in the future will help to achieve speeds compatible with live cell imaging.

Figure 9 summarises the results of the macro- and microscale BM measurements within cellular and acellular hydrogel models. The microscale measurements taken within the neuronal cell area exhibit BFSs higher on average than the BFSs of acellular gels and the hydrogel component of cellular gels. This is expected, since the macroscale measurements are taken using a low NA objective lens and thus present information on the mechanical properties averaged across a relatively large volume (approx. 200 μm^3^) of the aggregate material (cells and gel), hence lack the spatial resolution required to resolve the cells. Our microscale imaging setup has sufficient resolution (focal volume of approx. 3 μm^3^) to resolve individual cells. Local microscale mechanical measurements demonstrated that neuronal cells grow, multiply and form spheroid-type structures, with BFSs significantly higher (5.5%) than that of the surrounding hydrogel over 7 days of the experiment.

**Fig 9.**
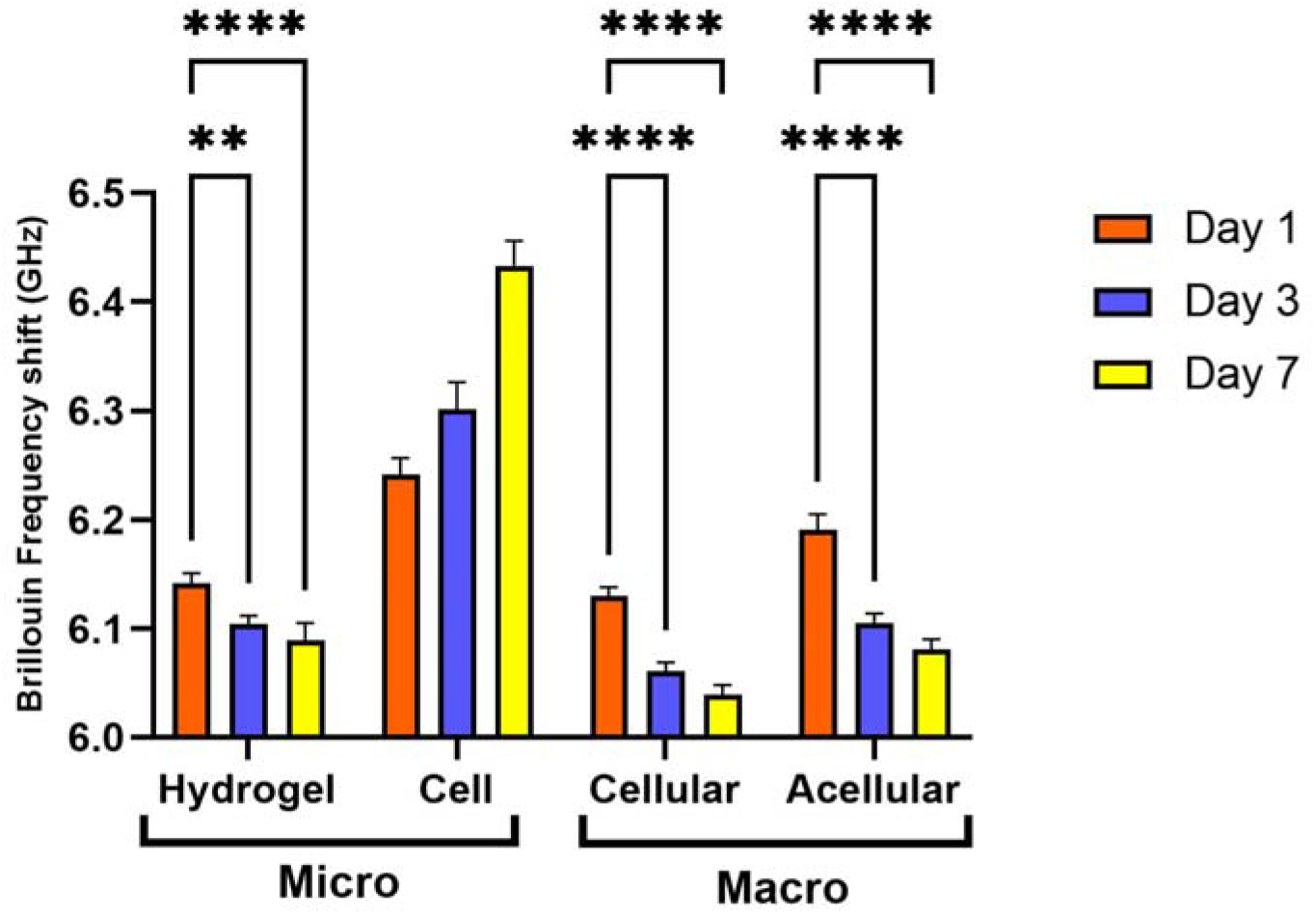
BFS measurements of the bioprinted acellular and cellular GelMA hydrogels embedded with NG108-15 neuronal cells on both the macro and microscale at concentrations of 2.5% (w/v). Error bars indicate 95% confidence interval; lines indicate statistical difference determined by two way ANOVA. Significance levels: **P <0.005, ****P <0.0001, n = 3.

Based on BM capabilities to resolve microscale viscoelastic properties in 3D models over time, we believe that this technology can help to customise physiological stiffness and mechanical support for CNS cells in 3D in vitro injury models. Injury to the CNS usually causes glial scar formation, which is significantly softer (∼50 Pa) than healthy CNS tissues (∼1 kPa) [1]. Following glial scar formation, astrocytes have been found to change their phenotype from naive astrocytes to reactive astrocytes [41]. The phenotypic changes of astrocytes are affected by multiple microenvironmental cues, one of which is matrix stiffness [42]. Understanding the changes in matrix stiffness and being able to accurately measure these in a non-destructive way may allow us to develop strategies to alter the stiffness of the matrix and as a result, control the reactivity of astrocytes in complex multicellular in vitro models. This may allow enhanced neuronal regeneration in vitro, giving rise to neuroregenerative strategies for spinal cord injury patients in the future.

## 5. Conclusions

Overall, we have demonstrated that Brillouin Microscopy can be applied to resolve macro- and microscale mechanical properties in cellular and acellular 3D bioprinted neuronal models over time. By using macroscale measurements, the BFS of hydrogels in a cell-free environment was found to change over the incubation period possibly due to gel degradation. Loading hydrogel constructs with cells accelerated these changes which we postulate occur through cell-matrix interactions. Furthermore, it was demonstrated BFS changes in cellular hydrogels were related to both solid fraction and incubation time, where the BFS variations of the hydrogels with lower solid fractions were highly significant compared to those of the higher solid fractions. Therefore, it is important to consider the mechanical properties as a function of incubation time rather than as a static property. In addition, by assessing cellular hydrogels on the microscale, the mechanical properties of hydrogels were measured within and surrounding the cellular spheroids, we demonstrated that the spheroid-type structures became stiffer over time. Overall, BM was found to be a useful tool for characterisation and deeper understanding of how the extracellular matrix in neural models change over time. This in situ mechanical characterisation method may be valuable for future studies and development of potential therapeutic approaches for neuronal regeneration and controlling glial scar formation following spinal cord injury.

## Credit authorship contribution statement

**Maryam Alsadat Rad**: Conducting a research and investigation process, design of methodology, Formal analysis, Visualization, Writing – original draft. **Hadi Mahmodi**: Formal analysis, Brillouin microscopy data collection, Writing-review & editing. **Elysse C.Filipe**: Rheology/compression analysis, Development and design of methodology. **Thomas R. Cox**: Rheology/compression analysis, Investigation/Methodology/Writing-Review & Editing. **Irina Kabakova**: Supervised Brillouin microscopy experiments, data analysis and interpretation and assisted in manuscript drafting and editing. **Joanne L. Tipper**: Supervised the findings of this work, Data analysis and interpretation and assisted in manuscript drafting and editing.

## Acknowledgments

This work was supported by the award of a scholarship to Maryam Alsadat Rad from the Faculty of Engineering and Information Technology (FEIT), UTS. TRC and ECF are supported by the NHMRC and CCNSW.

## Appendix A. Supplementary data

